# Practice-dependent refinement of motor execution is retained and broadly transferable but constrained by movement direction

**DOI:** 10.64898/2026.03.20.713284

**Authors:** Raphael Q. Gastrock, Setayesh Nezakatiolfati, Andrew King, Denise Y. P. Henriques

## Abstract

Practice enhances motor acuity, enabling movement execution with greater speed and accuracy. However, the learning principles underlying improvements in speed, accuracy, and efficiency remain less understood than those supporting motor skill acquisition and adaptation. Here, we examined motor execution in a skill-based practice task to characterize learning, retention, and generalization of motor acuity. Using a gamified two-dimensional racing task, right-handed participants controlled a stylus-driven car along a curved track as quickly and accurately as possible. Across two studies (N = 83 total, 54 females), participants completed 300 training laps on Session 1 and returned for Session 2 to assess retention and generalization to novel track configurations: one with altered spatial configuration (rotated track) and one requiring movement in the opposite direction of training (reverse track). Movement speed improved rapidly and showed robust, though incomplete, retention across sessions. Speed improvements generalized substantially to both novel tracks. Accuracy was high at training onset and showed strong retention. However, we do not observe offline gains between sessions. Notably, accuracy declined transiently for the novel track configurations, suggesting interference from prior training. Movement efficiency, indexed by path length, was retained and generalized to the rotated track. However, reversing movement direction impaired efficiency, revealing a movement direction effect. This effect persisted when training direction was reversed in a second study, with counterclockwise movements remaining slower and less efficient than clockwise movements. These findings show that practice produces durable and broadly transferable motor execution improvements, while inherent movement direction biases constrain how improvements generalize across contexts.

**New & Noteworthy:** The learning principles underlying improvements in motor acuity remain less well understood than those governing other forms of motor learning. Prior work suggests that motor execution improvements show limited generalization. In contrast, the present findings demonstrate that execution-based practice can produce robust, transferable gains, while also revealing a key constraint: inherent movement direction biases that limit generalization. By characterizing learning, retention, and generalization, this work provides new insight into how motor acuity improvements compare with skill acquisition and adaptation.

## Introduction

Humans are remarkable in their ability not only to acquire new motor skills or adapt well-known movements to perturbations, but also to refine movement execution through continued practice. For example, although an individual may learn the appropriate control policy to throw a ball toward a target, extensive practice is required to achieve the level of execution seen in a professional pitcher. Accordingly, most motor learning research on skill acquisition and adaptation has emphasized how appropriate actions are selected (1,2). Beyond action selection, however, many everyday behaviors benefit from practice-driven improvements in motor acuity, enabling movements to be executed with greater speed and accuracy (3–6). The learning mechanisms underlying these improvements in motor execution remain relatively understudied, and it is unclear how the processes supporting motor skill acquisition and adaptation relate to those governing practice-based refinements in motor execution. Here, we investigate continued improvements in motor execution during skill-based practice, using established measures from acquisition and adaptation paradigms to characterize the learning, retention, and generalization of motor acuity improvements.

Previous skill-based practice tasks have shown that performance variability reliably decreases with practice. One example is the arc-pointing task (4–6), in which participants use their wrist to perform point-to-point movements that guide a cursor from a start position to a target along a curved arc while remaining within a predefined pathway. Improvements in this task are reflected in reduced trajectory variability across practice. Changes in speed and accuracy further reveal a characteristic speed-accuracy trade-off. That is, although longer movement times initially produce greater accuracy within the arc, accuracy continues to improve across multiple days of training (4).

Similar patterns have been reported in related track-tracing tasks (7,8), which likewise demonstrate systematic changes in speed and accuracy with practice. Notably, however, both the arc-pointing and track-tracing paradigms impose temporal constraints on movements to assess speed-accuracy trade-offs. In contrast, Petersen et al. (9) examined performance under conditions in which participants were free to move as quickly and accurately as possible, without visual feedback, instead relying on memory of the correct path. Building on these previous works, we introduce a custom two-dimensional racing task in which participants were instructed to maximize both speed and accuracy. This design allows us to examine practice-related improvements in speed, accuracy, and performance variability without imposing constraints on movement speed or visual feedback.

Improvements in motor execution have distinct underlying neural processes compared to those that contribute to skill acquisition and adaptation. Previous research in both non-human and human primates have shown that practice-related improvements are primarily supported by motor cortical areas (5,1), whereas skill acquisition and adaptation recruit other areas including the prefrontal cortex, basal ganglia, and cerebellum (1,10–12). Nevertheless, practice-based improvements reflect similarities in learning processes when compared to those that are known from skill acquisition and adaptation. One example is a similarity between the interplay of explicit and implicit processes, an interaction that has been well characterized in studies of motor skill acquisition and adaptation (13–20). Mastering a motor skill typically requires extended practice, where expertise development is often attributed to deliberate practice, a cognitively demanding form of training involving structured activities aimed at improving performance rather than simple task repetition (21,22). Recent computational work, however, challenges this view, suggesting that more explicit or cognitive involvement in motor learning may be largely confined to early stages of acquisition, with practice-based performance improvements relying less on cognitive processes (2). Here, we focus on other behavioral patterns known from acquisition and adaptation tasks, including how learning is retained over time and its generalization to novel contexts.

Skill acquisition tasks show robust patterns of retention, particularly when compared with the more limited retention observed in adaptation tasks (23,24,20). One hallmark of such robust retention is the presence of offline gains, where performance at the start of a second session exceeds the asymptotic performance reached during the initial training session (23). Comparable retention effects have been reported in skill-based practice tasks, in which accuracy continues to improve across multiple days of training (4–8). With respect to generalization, we have previously shown substantial transfer of performance to novel conditions in a skill acquisition task (20). In skill-based practice paradigms, such as the arc-pointing task, training at a specific movement speed supports transfer to other speed ranges (4). However, generalization in these tasks appears to be constrained by handedness and seems specific to the trained movement trajectory (7,6). Together, these findings suggest that generalization of motor execution improvements to novel conditions may be limited. Here, we specifically examine retention and generalization of performance in our racing task, while separately assessing generalization based on trained movement direction and track configuration.

We developed a gamified two-dimensional racing task in which participants used a stylus on a digitizing tablet to control a race car displayed on a monitor. Their goal was to move through the track as quickly and accurately as possible, with off-track movements penalized. The experiment spanned two days. On Day 1, participants completed 300 training laps on a single track. On Day 2, we first assessed retention on the trained track, then tested generalization by presenting the same track in a new orientation and another configuration where participants had to move through the track in the reverse direction. We hypothesize robust retention and substantial transfer to these novel conditions, consistent with patterns observed in skill acquisition. Although training required clockwise movements, performance declined when participants moved through the track in the reverse counterclockwise direction, revealing a movement direction effect. To investigate this further, we conducted a second study directly comparing performance for clockwise and counterclockwise movements. We hypothesize that this effect reflects inherent movement biases rather than interference from exposure to the novel track configurations.

## Methods

### Participants

For Study 1, 45 right-handed adults (28 female; *M*_age_ = 21.33, *SD*_age_ = 4.47) participated in Session 1, and 43 participants (26 female) returned for Session 2. For Study 2, 38 right-handed adults (26 female; *M*_age_ = 20.68, *SD*_age_ = 2.55) completed Session 1, with 36 participants (25 female) returning for Session 2. All participants reported normal or corrected-to-normal vision and had no known neurological disorders, except for two participants in Study 1 and one participant in Study 2 that reported having a neurological condition (e.g., ADHD or OCD). Inspection of their task performance revealed no qualitative differences relative to the remaining participants, and we therefore included their data in all analyses. We based our recruitment sample from previous similar studies (4,5) and ensured sufficient representation of participants trained in each of the trained track orientations. All participants provided written informed consent prior to participation and received course credit. Experimental procedures complied with institutional and international guidelines and were approved by York University’s Human Participants Review Committee.

### Experimental set-up

#### Apparatus

Participants sat on a height-adjustable chair with a digitizing tablet (Wacom Intuos Pro, 16.8” x 11.2” x 0.3”, resolution resampled to 1920 x 1080 pixels at 60 Hz) directly in front of them and a vertically mounted monitor (Dell, 24” P2422H) positioned 30.5 cm ahead of the tablet (Fig. 1A). Using their right hand, they moved a digitizing stylus across the tablet surface while viewing task-related stimuli on the monitor. We instructed participants to sit comfortably to avoid physical strain during the experiment.

**Figure 1.**
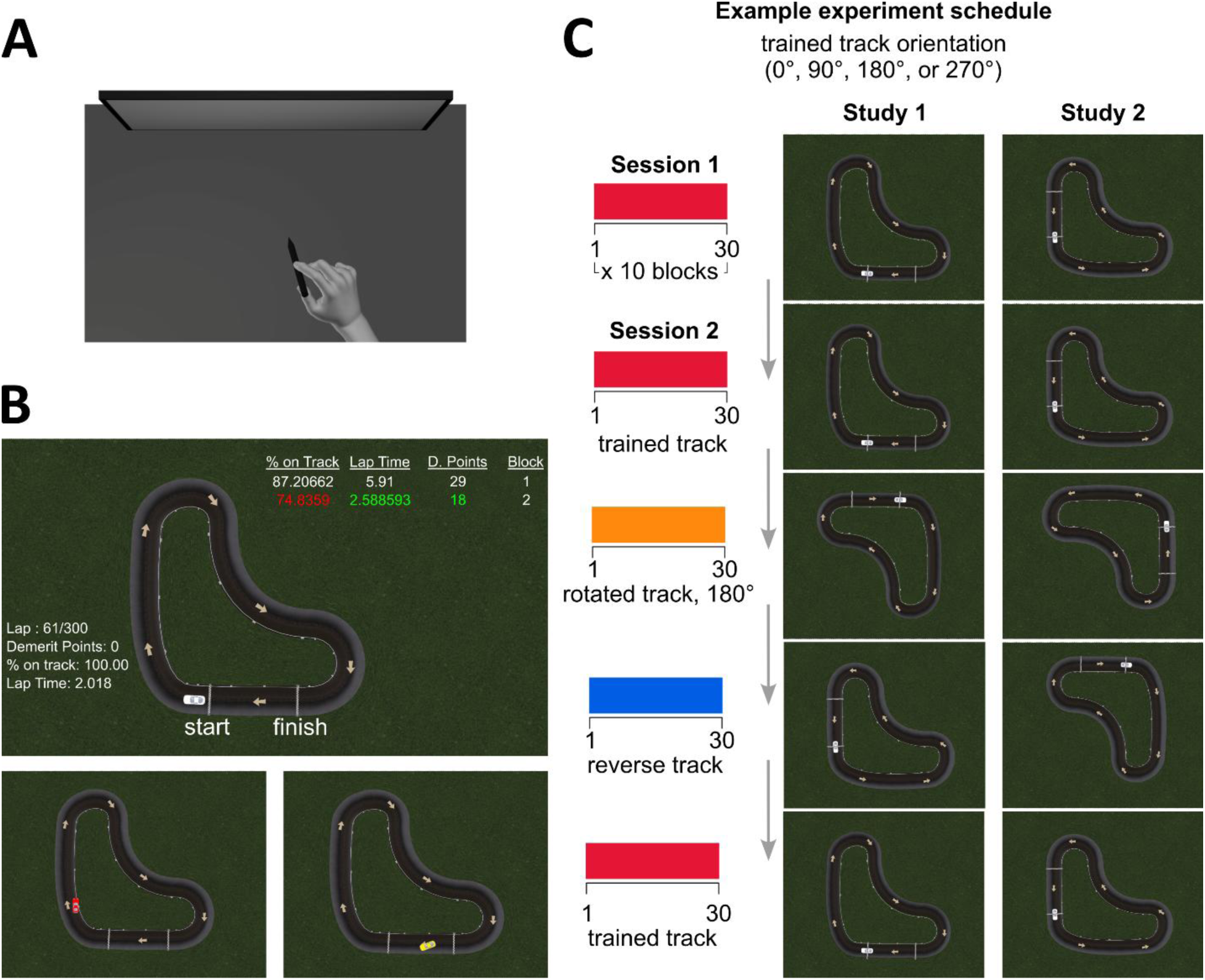
Experimental set-up and design. (**A**) Participants used their right hand to control a stylus on a digitizing tablet while viewing the task on a vertically mounted monitor. (**B, top**) Two-dimensional racing task viewed from a bird’s-eye perspective. Participants guided an on-screen cursor (white car) from start to finish in the direction indicated by arrows. The start and finish lines are labelled here for reference, but these labels were not visible to participants during the experiment. After each lap, feedback displayed lap number, demerit points (track boundary violations), percentage on track, and lap time. After each block of 30 laps, summary feedback showed average percentage on track, average lap time, and total demerit points, with improvements relative to the previous block shown in green and red otherwise. (**B, bottom**) The car turned red when outside the track boundaries and yellow within the pit zone, where participants could rest between laps. (**C**) To create distinct training contexts while preserving identical track geometry, participants were assigned to one of four track orientations (0°, 90°, 180°, or 270° in polar coordinates). For both studies, Session 1 consisted of 300 training laps on the assigned track. In Study 1, Session 2 included 30 laps on the trained track, followed by 30 laps on the track rotated by 180° (rotated track). This rotated track was then further rotated by 90° and flipped along either the horizontal (0° and 180°) or vertical (90° and 270°) axis, after which participants completed 30 laps in the reverse direction (reverse track). Finally, participants completed another 30 laps on the trained track. Study 2 followed a similar structure for Session 2 but did not further alter the reverse track, allowing assessment of performance in the reverse direction alone. Movements were in the clockwise direction for Study 1 and counterclockwise for Study 2.

#### Stimuli

We developed a custom two-dimensional racing task using Unity (Fig. 1B; version 2021.3.2f1, Unity Technologies). In Unity, track spatial dimensions were scaled differently along the horizontal (62 px = 1 m) and vertical (57 px = 1 m) axes of the workspace, and were subsequently mapped onto both the display screen (36 px/cm) and the tablet. Accordingly, the dimensional conversions were as follows: horizontal (1 m in Unity = 1.72 cm on the screen = 1.01 cm on the tablet); vertical (1 m in Unity = 1.58 cm on the screen = 1.15 cm on the tablet). To maintain consistency, all our reported dimensions and movement distance measures are expressed in screen-based centimeters.

Participants viewed the track, positioned at the center of the screen, from a bird’s-eye perspective (Fig. 1B; overall track width = 17 cm, height = 17.3 cm). Stylus movements controlled an on-screen cursor displayed as a white car (width = 0.7 cm, length = 1.6 cm). We instructed participants that their goal for the task was to navigate the car from the start to the finish line in the direction indicated by arrows on the track, as quickly as possible, while maintaining accuracy by keeping the car within the track boundaries (2 cm wide). We implemented several task features to encourage participants’ continuous monitoring of performance and sustained attempts to improve speed and accuracy. First, when the car crossed the track boundaries, it turned red and an auditory beep was triggered (Fig. 1B, bottom left), signaling participants to return the car to the track. Upon re-entry, the car reverted to its default white color. Second, each boundary violation resulted in a demerit point, indexing the number of times a participant exceeded the track boundaries within a lap. Third, at the end of each lap, participants received immediate performance feedback displayed on the left side of the screen (Fig. 1B, top), including lap number, demerit points, percent on track (defined as the percentage of time the car remained within the track boundaries during the lap), and lap time. Fourth, after each block of 30 trials, participants were shown their average lap time, average percent on track, and total demerit points for that block (Fig. 1B, top). From the second block onward, these summary measures were displayed in green if performance improved relative to the previous block and in red otherwise. Finally, to minimize fatigue, participants were encouraged to take breaks between laps. During breaks, participants positioned the car in the pit zone (Fig. 1B, bottom right; the area between the finish and start lines), where the car turned yellow and the timer was paused.

Participants moved the car along a curved, closed race track with a fixed geometry resembling an elbow-shaped loop (Fig. 1B). To create distinct training contexts while preserving identical track geometry, we rotated the same track into one of four orientations (0°, 90°, 180°, or 270° in polar coordinates; Fig. 1C). Track orientation was counterbalanced across participants. The trained orientation also determined the corresponding track conditions experienced by each participant in Session 2 (see Procedure). However, we observed no qualitative differences in performance across orientations (see R notebook: 10.17605/OSF.IO/W5D4U). Thus, we combined and analyzed data from all track orientations together.

#### Study 1 Procedure

We instructed participants that the goal of the task was to use the stylus on the digitizing tablet to guide the car smoothly around the track, in the direction indicated with arrows, as quickly and accurately as possible. During Session 1, participants trained for 300 laps on their randomly assigned track orientation (Fig. 1C). Each lap began when the car crossed the start line from the pit zone and ended when it crossed the finish line and re-entered the pit zone. Regardless of track orientation, all laps in Session 1 required a clockwise movement around the track.

In Session 2, participants returned after a minimum of one day (inter-session interval: *M*_day_ = 1.02, *SD*_day_ = 0.15), and we assessed for retention of learning from Session 1 and performance generalization to altered track conditions. We evaluated retention by having participants immediately complete 30 laps on their trained track orientation at the start of Session 2 (Fig. 1C, trained track). We then assessed for generalization across different novel track conditions. First, participants completed 30 laps on a track rotated 180° relative to their trained orientation (Fig. 1C, rotated track). Next, the track was further rotated by 90° and flipped along either the horizontal (for 0° and 180° track orientations) or vertical (for 90° and 270° track orientations) axis, such that the orientation of the track resembled the trained track orientation but with different start and finish positions. We then instructed participants to move through this track in the reverse (counterclockwise) direction for 30 laps (Fig. 1C, reverse track). Finally, participants completed an additional 30 laps on the original trained track orientation (Fig. 1C, trained track). Session 2 ended once all 120 laps were completed.

#### Study 2 Procedure

Study 1 revealed an unexpected movement direction effect, whereby reverse (counterclockwise) movements around the track impaired movement speed and trajectory quality, relative to trained clockwise movements (see Results). Accordingly, in Study 2 we replicated the experiment in an independent sample using the same apparatus and procedures, while reversing the trained movement direction (Fig. 1C, Study 2). Specifically, participants trained moving through the track in the counterclockwise direction during Session 1, and all trained track and rotated track laps in Session 2 also required counterclockwise movement. For the reverse track, we did not further alter the track orientation from the preceding rotated block. Instead, participants were instructed to move clockwise, allowing us to isolate performance assessment in the reverse direction. Participants completed a total of 300 laps during Session 1, and 120 laps during Session 2. Participants returned for Session 2 after a minimum of one day (inter-session interval: *M*_day_ = 1.17, *SD*_day_ = 0.45).

### Data analysis

For both studies, each block of 30 laps was divided into five sets of six laps. We quantified learning, retention, and generalization using the first and last sets of laps across selected blocks in the two sessions. At the start of each experiment, the car spawned on the starting line. However, placing the stylus on the digitizing tablet repositioned the car to the stylus location, occasionally triggering unintended lap starts. In Session 2, the starting line and car spawn location changed at the onset of each novel track configuration (Fig. 1C). Consequently, performance on the first lap of Session 1 and the first lap of every block in Session 2 was confounded by these inadvertent starts. We therefore excluded the first lap of every block from all analyses, retaining five laps in each first lap set and six laps in each last lap set. We first analyzed performance changes within each study independently and then conducted direct comparisons between studies. We report Bayesian statistics to show evidence in support of the alternative hypothesis over the null hypothesis (BF_10_ values). We include Bayes Factors (BF) for the comparisons we conduct and inclusion Bayes Factors (BF_incl_) when assessing main and interaction effects of predictors. Reported Bayes factors reflect the ratio of how likely the alternative hypothesis is (i.e., there is a difference) over how likely the null hypothesis is (i.e., there is equivalence), given a non-informative prior and the data. With BF_10_ = 1 both are equally likely. Within the interval 1/3 to 3 (either hypothesis is up to 3 times more likely than the other) there is only anecdotal evidence, and thus no real effect (25,26). However, BF_10_ values outside this interval are considered evidence in favor of either the alternative or null hypothesis. Note that in contrast to classic null-hypothesis significance testing (NHST), Bayesian statistics allow confirming a null hypothesis. We also report more detailed results from both Bayesian and corresponding frequentist tests in our accompanying R notebook (10.17605/OSF.IO/W5D4U). All data preprocessing and analyses were conducted in R version 4.2.2 (27).

We quantified performance changes across training sessions using different dependent measures: movement speed (lap time), accuracy (percentage on track), and movement efficiency (path length).

#### Lap time

We defined lap time as the time elapsed between the first sample after the car crosses the start line and the final sample when the car crossed the finish line. To assess retention, we compared performance on the first and last set of laps in Session 1 with the first set in Session 2. To assess generalization, we compared performance on the first set of laps for each of the two novel track orientations in Session 2 (i.e., rotated and reverse tracks) with performance on the first and last set of laps in Session 1. Since we counterbalanced the track orientation across participants, initial performance on a novel track can be directly compared with initial performance of other participants when first exposed to that same track orientation. Thus, any improvement on the novel tracks reflects transfer of skill beyond the specific trained orientation, rather than familiarity with that particular track.

#### Percentage on track

We defined accuracy as the proportion of time (expressed as a percentage of total lap time) that the car remained within the track boundaries during each lap. We compared percentage on track across different sets of laps in both sessions, to assess retention and generalization patterns using the same analysis steps outlined for lap times.

#### Path length

We assessed movement efficiency using path length (PL), defined as the total distance travelled between lap start and finish times, calculated from the x and y coordinates of all recorded points on the movement trajectory. The midline of the track served as a reference, with a total length of 49.68 cm from the start to the finish line. In addition to total PLs, we separately calculated path length for movements made within the track boundaries (PL inside track) to assess changes in movement efficiency independent of accuracy. As with the other dependent measures, PL and PL inside track were compared across different sets of laps in both sessions to assess retention and generalization.

## Data availability

Data and analyses scripts are available on Open Science Framework (10.17605/OSF.IO/W5D4U).

## Results

Since we measured the same dependent variables within each study, we first report performance improvements separately for each study. We assessed performance using one-way repeated-measures Bayesian ANOVAs with lap set as the within-subjects factor for each of the dependent measures: lap time, percentage on track, and path length. We then assessed planned comparisons between specific lap sets using Bayesian t-tests. While we present trial-by-trial data for each dependent measure (Fig. 2-5), we also show the relationship between speed and accuracy improvements in figure 3. Finally, we report direct comparisons between the two studies as a separate section at the end.

**Figure 2.**
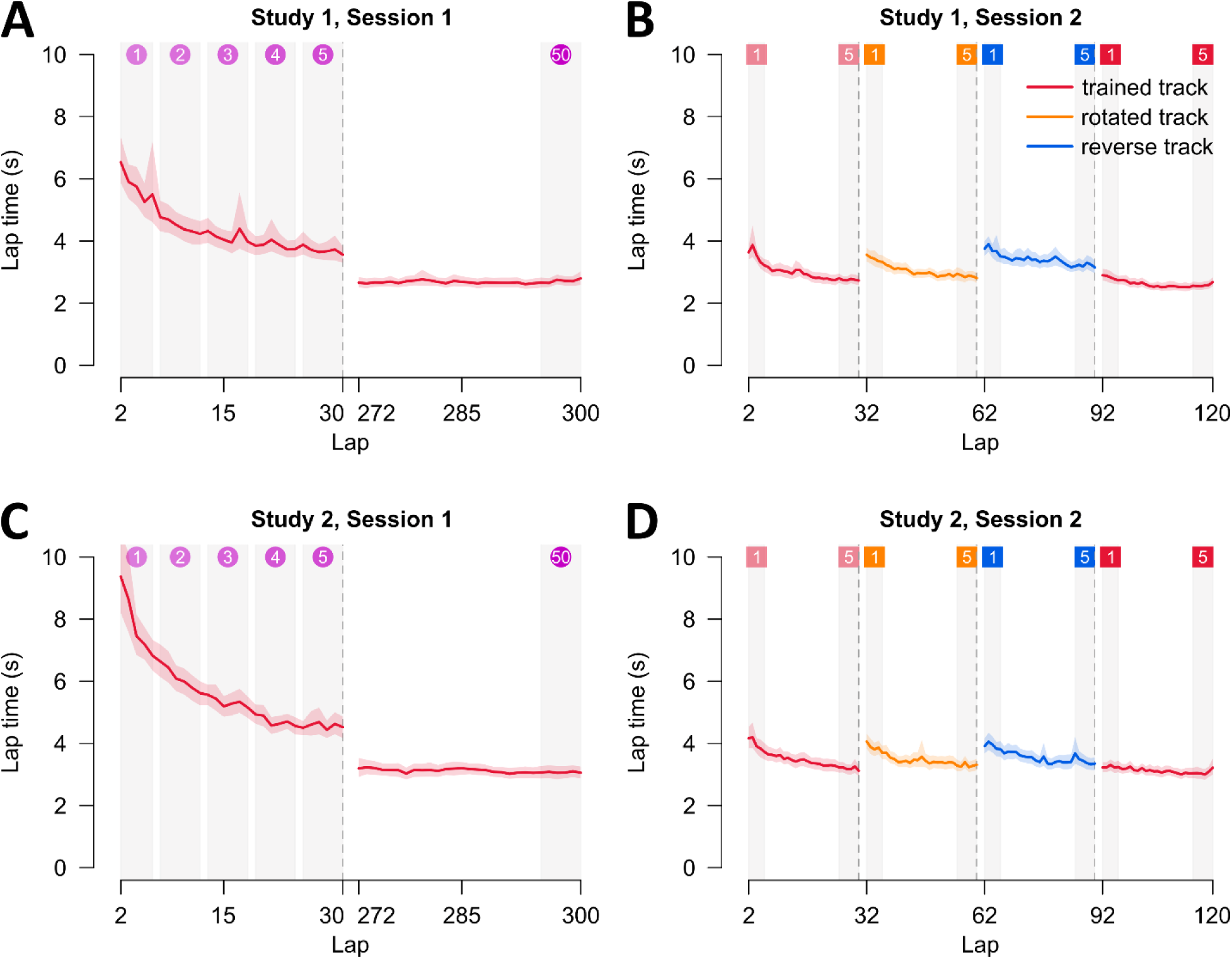
Lap times across sessions for both studies. Lap time is the total time elapsed between the first sample after the car crosses the start line and the final sample when the car crossed the finish line. (**A**) For Study 1, we include the first and last blocks of training in Session 1. (**B**) Session 2 includes every block with the trained track, rotated track, reverse track, and final trained track block. (**C-D**) Similar to panels A-C but with data from Study 2. For panels A-D, mean lap time is presented as solid lines, and shaded regions correspond to 95% confidence intervals. The numbered lap sets at the top of each panel correspond to the sets used in the statistical comparisons conducted.

**Figure 3.**
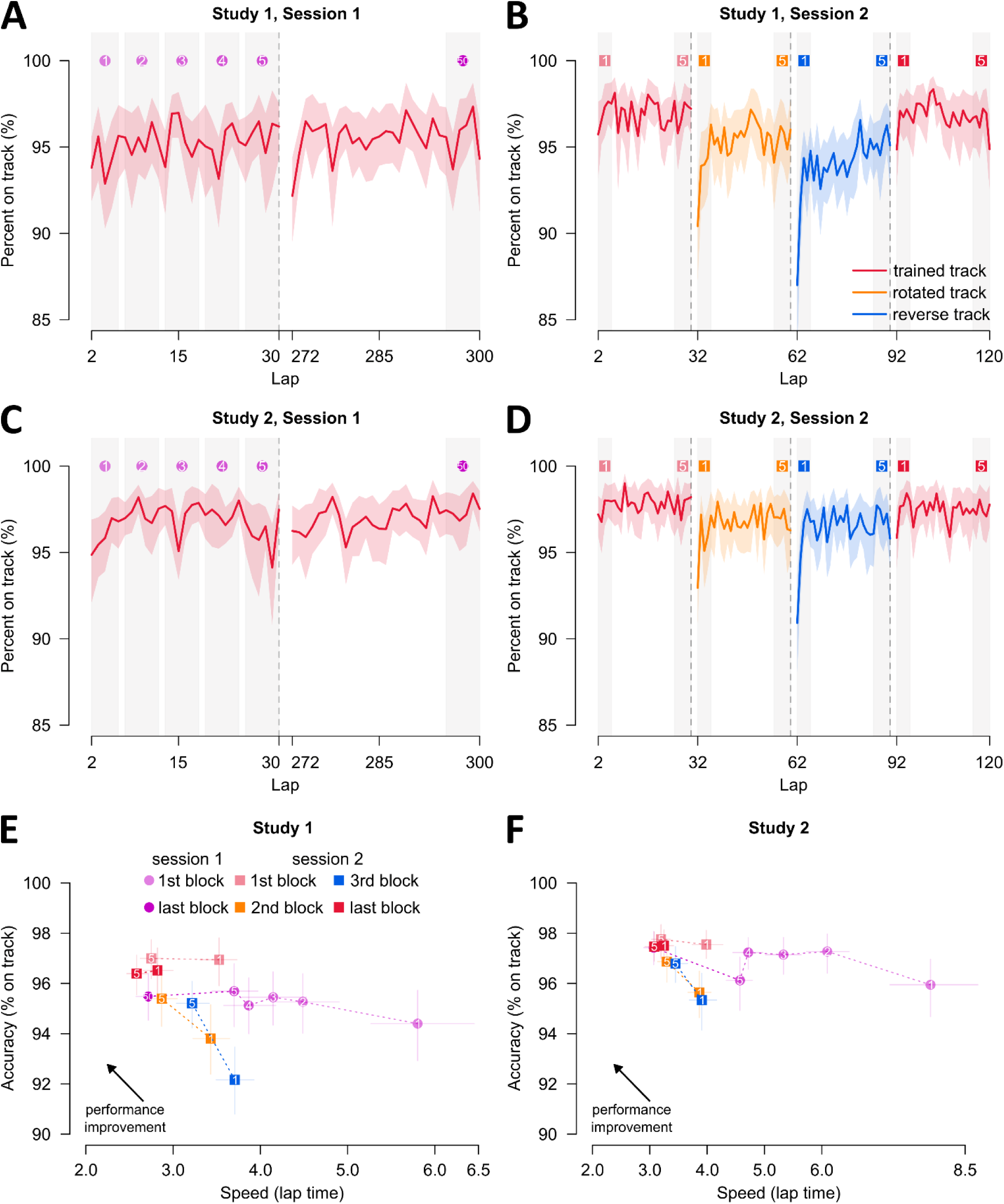
Accuracy across sessions for both studies. Accuracy is defined as the proportion of time (expressed as a percentage of total lap time) that the car remained within the track boundaries during each lap. (**A**) For Study 1, we include the first and last blocks of training in Session 1. (**B**) Session 2 includes every block with the trained track, rotated track, reverse track, and final trained track block. (**C-D**) Similar to panels A-C but with data from Study 2. For panels A-D, mean percentage on track is presented as solid lines, and shaded regions correspond to 95% confidence intervals. (**E**) Speed and accuracy improvements for Study 1. We divided each block of 30 laps into five sets of six laps. In Session 1, the first block is shown as light purple circles, separated into 5 sets (numbered 1 to 5). The last block is shown as a dark purple circle, and only includes the last trial set (numbered 50). In Session 2, we only show the first and last set of every block (numbered 1 and 5), where the first trained track block is shown as light red squares, the rotated track as orange squares, the reverse track as blue squares, and the final trained track block as dark red squares. These numbered sets correspond to the lap sets used in the statistical comparisons, and the lap-by-lap data are labeled accordingly in panels A–D. (**F**) Similar to panel E but with data from Study 2. For panels E-F, solid shapes represent mean values, and solid lines correspond to 95% confidence intervals for either dependent variable.

### Improvements in speed show partial but robust retention and generalization to novel track configurations

We first assessed improvements in speed, quantified by lap time. In both studies, lap times decreased steadily over Session 1 and approached an asymptote, plateauing at an average of 2.68 s in Study 1 and 3.09 s in Study 2 (Fig. 2A, 2C). Comparisons of performance on the trained track across the first and last sets of Session 1 and the first set of Session 2 revealed strong evidence for an effect of set in both studies (Study 1: BFincl > 10²⁰; Study 2: BFincl > 10²⁵). This effect was partly driven by robust within-session improvements during Session 1, with lap times decreasing by an average of 3.04 s in Study 1 (Fig. 2A; BF > 10¹⁰) and 4.74 s in Study 2 (Fig. 2C; BF > 2 × 10¹¹). In Session 2, participants began with substantially faster lap times than at the start of Session 1, improving by an average of 2.20 s in Study 1 (Fig. 2B; BF > 2 × 10⁷) and 3.85 s in Study 2 (Fig. 2D; BF > 10⁹). However, initial performance in Session 2 remained slower than the final set of Session 1 by an average of 0.84 s in Study 1 (BF > 10⁸) and 0.89 s in Study 2 (BF > 2 × 10⁵). Taken together, these results indicate rapid learning of the racing task, accompanied with substantial but incomplete retention of lap time improvements across sessions.

We next examined generalization of performance to novel track configurations. For the rotated track, we compared lap times from the first set of laps on this novel track in Session 2 (orange curves in Fig. 2B, 2D) with the first and last sets of laps on the original, trained track in Session 1. This analysis revealed strong evidence for an effect of set in both studies (Study 1: BFincl > 4 × 10¹⁹; Study 2: BFincl > 3 × 10²⁶). Participants began the rotated track in Session 2 with faster lap times than at the start of Session 1, showing average improvements of 2.29 s in Study 1 (Fig. 2B; BF > 5 × 10⁶) and 3.96 s in Study 2 (Fig. 2D; BF > 3 × 10¹⁰). However, initial performance on the rotated track remained slower than the final set of Session 1 by 0.75 s in Study 1 (BF > 10⁶) and 0.78 s in Study 2 (BF > 4 × 10⁶). We conducted an analogous analysis for the reverse track (blue curves in Fig. 2B, 2D). Again, there was strong evidence for an effect of set in both studies (Study 1: BFincl > 4 × 10¹⁹; Study 2: BFincl > 4 × 10²⁵). Initial lap times on the reverse track in Session 2 were faster than initial lap times in Session 1 by an average of 2.01 s in Study 1 (Fig. 2B; BF > 2 × 10⁶) and 3.91 s in Study 2 (Fig. 2D; BF > 3 × 10⁹). As with the rotated track, initial performance on the reverse track remained slower than the final set of Session 1, by 1.03 s in Study 1 (BF > 5 × 10¹⁰) and 0.83 s in Study 2 (BF > 8 × 10⁴). Together, these findings indicate substantial, though incomplete, generalization of lap time improvements to both rotated and reverse track configurations.

Finally, we examined whether experience with the novel track configurations affected performance upon re-encountering the original trained track. To this end, within Session 2 we compared lap times from the last set of laps in the first trained track block with those from the first set of laps in the final trained track block (trained track blocks in Fig. 2B, 2D). In both studies, lap times did not differ between these two sets (Study 1: BF = 0.24; Study 2: BF = 0.22), indicating no changes in performance. These results suggest that training on the novel track configurations does not interfere with performance on the original trained track.

### Accuracy improvements are retained, though performance is initially reduced on novel track configurations

We next assessed improvements in accuracy. In both studies, participants achieved high accuracy almost immediately, with the car remaining within the track boundaries for over 90% of each lap (Fig. 3). Accuracy changed only modestly across training. Although there was an overall effect of set in both studies (Study 1: BFincl = 5.83; Study 2: BFincl = 15.94), within-session improvements were minimal (Fig. 3A, 3C). In Study 1, there was no evidence for within-session improvement (Fig. 3A, 3E; BF = 0.28), whereas in Study 2 accuracy increased by 1.7% (Fig. 3C, 3F; BF = 5.34), indicating modest gains. These limited improvements likely reflect ceiling effects arising from the high initial accuracy. Consistent with this, the number of demerit points incurred when the car left the track was also low from the outset and showed little change across training (see R notebook: 10.17605/OSF.IO/W5D4U), declining modestly in Study 1 (on average 1.20 to 0.91 demerit points) and Study 2 (2.41 to 1.23 demerit points). Overall, participants performed near ceiling from the start, suggesting they prioritized staying on the track and directed any further improvement toward optimizing lap timing rather than accuracy.

When returning in Session 2, participants were moderately more accurate compared to the initial set of Session 1, by 2.5% in Study 1 (Fig. 3B, 3E; BF = 11.43) and 1.8% in Study 2 (Fig. 3D, 3F; BF = 3.96). Although accuracy was slightly higher in the first set of Session 2 than in the final set of Session 1 (average increase of 1.5% in Study 1; 0.1% in Study 2), our analyses did not show evidence for a difference between these sets (Study 1: BF = 2.10; Study 2: BF = 0.18). Taken together, these results indicate strong retention of accuracy across sessions, despite limited within-session improvement and no evidence for offline gains.

To assess generalization of accuracy to the novel track configurations, we compared accuracy from the first and last sets of laps on the trained track in Session 1 with the first set of laps on the rotated track in Session 2. As seen with the initial laps of orange curves in figures 3B and 3D, accuracy appears to be initially worse when the track was changed, compared to accuracy at the end of the first training session. We only find evidence for an effect of set for Study 2 (BFincl = 7.63), but none for Study 1 (BFincl = 0.38). The effect of set in Study 2 was driven by a clearer drop in accuracy (on average accuracy diminished by 1.9%) when participants began the rotated track in Session 2 compared to their last set on the original track in Session 1. This pattern was even more pronounced when participants encountered the second novel track configuration, the reverse track (blue curves in Fig. 3B, 3D). We again found evidence for an effect of set in both studies (Study 1: BFincl = 54.11; Study 2: BFincl = 9.78). In Study 1 (Fig. 3A-B, 3E), accuracy on the reverse track was lower than both the initial and final set of Session 1 by 2.2% and 3.2%, respectively (BF = 6.68 and 96.55). In Study 2 (Fig. 3C-D, 3F), a drop in accuracy of 2.2% was evident only relative to the final set in Session 1 (BF = 24.65). Taken together, these results show that accuracy is initially worse when participants transition to a novel track configuration, suggesting that training on the original track may transiently interfere with performance when first encountering a novel track configuration.

Finally, we again examined whether experience with the novel track configurations affected accuracy upon re-encountering the original trained track. As such, we compared percentage on track from the last set of laps in the first trained track block with those from the first set of laps in the final trained track block (trained track blocks in Fig. 3B, 3D). In both studies, accuracy did not differ between these two sets (Study 1: BF = 0.25; Study 2: BF = 0.21). Thus, training on the novel track configurations does not interfere with accuracy on the original trained track.

### Movement direction modulates movement efficiency

We next assessed improvements in movement efficiency, quantified by path length (PL). Using the track midline as a reference (49.68 cm; Fig. 4), participants in Study 1 consistently followed shorter, more efficient paths (Fig. 4A–4B), indicating that the optimal trajectory involves cutting inward during turns rather than strictly adhering to the midline. When comparing the first and last sets of laps on the trained track in Session 1 with the first set of laps in Session 2, we observed evidence for an effect of set (BF = 8.47). However, follow-up comparisons revealed no evidence for within-session improvement (BF = 2.41), no difference between the first sets of Sessions 1 and 2 (BF = 1.75), and no difference between the final set of Session 1 and the first set of Session 2 (BF = 0.20). Together, these results indicate stable movement efficiency on the trained track across sessions in Study 1.

**Figure 4.**
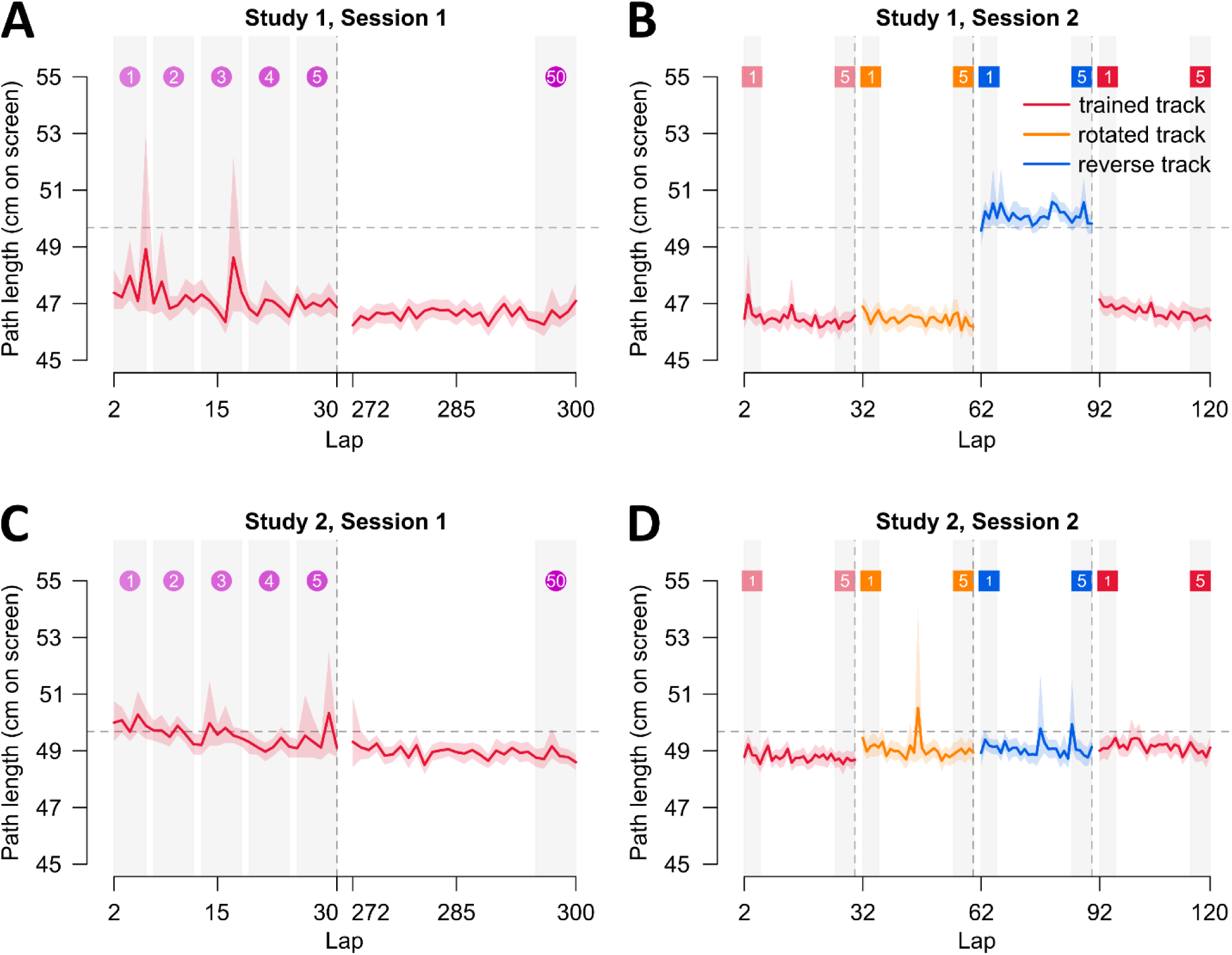
Path length across sessions for both studies. Path length is defined as the total distance travelled between lap start and finish times, calculated from the x and y coordinates of all recorded points on the movement trajectory. The midline of the track served as a reference, with a total length of 49.68 cm from the start to the finish line (horizontal dashed line in all panels). (**A**) For Study 1, we include the first and last blocks of training in Session 1. (**B**) Session 2 includes every block with the trained track, rotated track, reverse track, and final trained track block. (**C-D**) Path lengths during Sessions 1 and 2 for Study 2. For all panels, mean path length is presented as solid lines, and shaded regions correspond to 95% confidence intervals. The numbered lap sets at the top of each panel correspond to the sets used in the statistical comparisons conducted.

To assess generalization in Study 1, we compared path lengths from the first and last sets of laps on the trained track in Session 1 with the first set of laps on the rotated (orange curves in Fig. 4) and reverse (blue curves in Fig. 4) tracks in Session 2. These analyses revealed reduced movement efficiency only for the reverse track (Fig. 4B), with strong evidence for an effect of set (BFincl > 10¹³). This unexpected effect is clear from Fig. 4B, where the pathlengths were around 2-4 cm longer for the reversed track than for all other tracks. As mentioned earlier and will be described in more detail below, this finding motivated conducting study 2. More specifically, initial path lengths on the reverse track were longer than those in the first set of Session 1 by an average of 2.33 cm (BF > 10⁶) and longer than those in the final set of Session 1 by 3.49 cm (BF > 10¹⁶). Importantly, this effect persisted when the analysis was restricted to movements within the track boundaries (PL inside track; Fig. 5). Path lengths on the reverse track were longer compared to the trained track (Fig. 5B; BFincl = 23.89), exceeding those of the first set in Session 1 by an average of 1.2 cm (BF = 4.11) and the last set in Session 1 by 1.6 cm (BF = 97.59). In contrast, movement efficiency with the rotated track showed a different pattern. Initial path lengths inside the track were on average slightly shorter than those in Session 1 (1.2 cm shorter than the first set and 0.8 cm shorter than the last set; Fig. 5B), but there was no evidence for an effect of set (BF = 1.44). Together, these findings suggest that poorer performance on the rotated track is driven primarily by increased errors rather than inefficient trajectories, whereas the poorer performance on the reverse track reflects a reduction in movement efficiency that is independent of accuracy.

**Figure 5.**
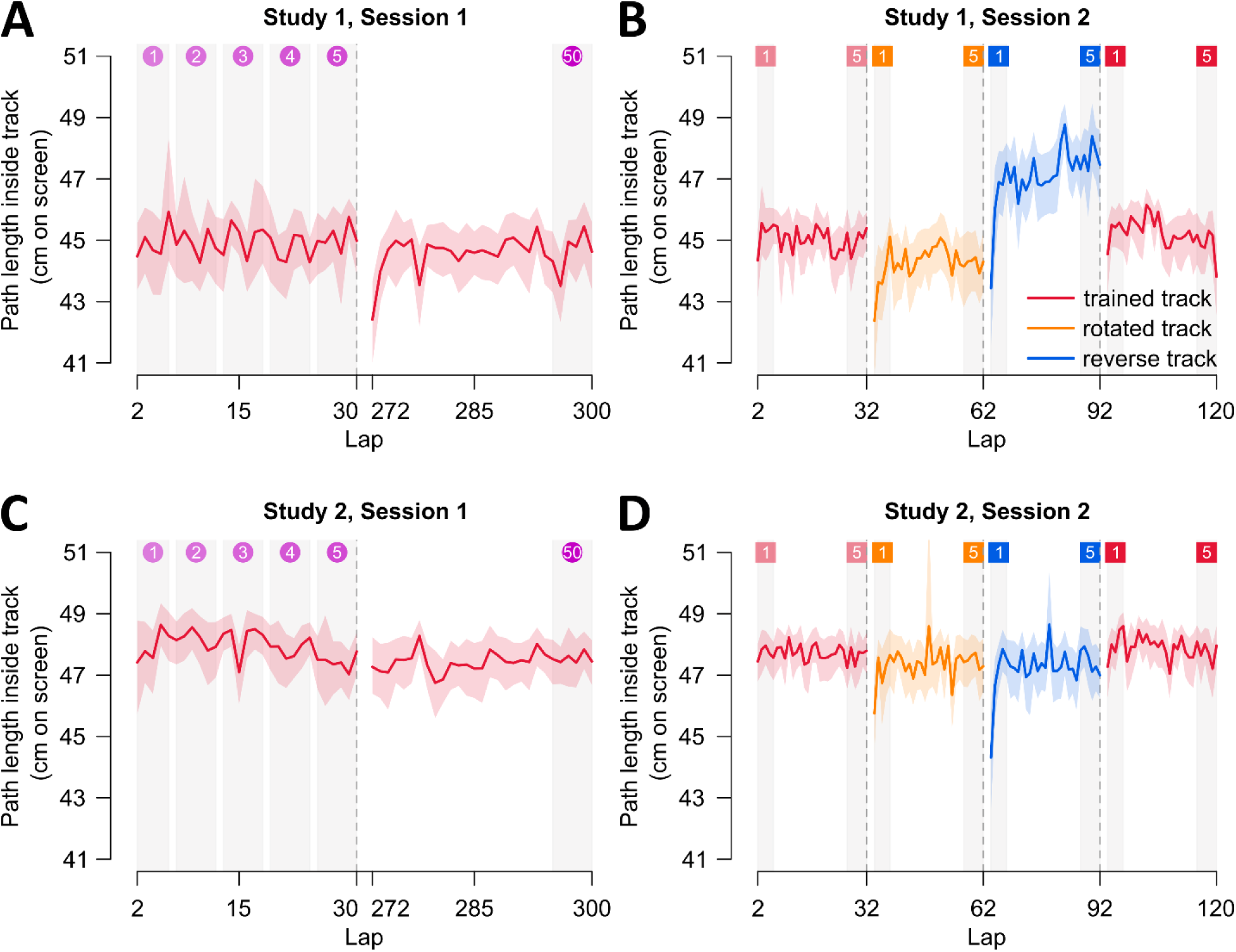
Path length inside track across sessions for both studies. Calculated path length for movements made within the track boundaries (PL inside track) to assess changes in movement efficiency independent of accuracy. (**A**) For Study 1, we include the first and last blocks of training in Session 1. (**B**) Session 2 includes every block with the trained track, rotated track, reverse track, and final trained track block. (**C-D**) Path lengths during Sessions 1 and 2 for Study 2. For all panels, mean path length is presented as solid lines, and shaded regions correspond to 95% confidence intervals. The numbered lap sets at the top of each panel correspond to the sets used in the statistical comparisons conducted.

Given the unexpected reduction in movement efficiency in the reverse track, this motivated Study 2 where we simply changed the trained driving direction. In Study 2, we also see that path lengths were several centimeters longer when participants had to move the car along the track in the counterclockwise direction, such that path lengths were closer to the track midline reference of 49.68 cm (Fig. 4C–4D). When comparing the first and last sets of laps on the trained track in Session 1 with the first set of laps in Session 2, we find strong evidence for an effect of set (BFincl > 3 × 10^8^). We find that this effect is driven by within-session improvements, where path lengths decreased by an average of 1.13 cm within Session 1 (Fig. 4C; BF = 9177.72). Moreover, initial path lengths in Session 2 are on average 1.11 cm shorter than initial path lengths in Session 1 (Fig. 4C-4D; BF > 7 × 10^4^) and do not differ from the final path lengths in Session 1 (BF = 0.18). Thus, in Study 2, we find that movement efficiency improves with training and is robustly retained across sessions.

We then assessed generalization in Study 2 by comparing path lengths from the first and last sets of laps on the trained track in Session 1 with the first set of laps on the novel track configurations (rotated and reverse tracks). For the rotated track, we find a main effect of set (BFincl > 10^6^). Initial path lengths in the rotated track are on average 0.77 cm shorter than initial path lengths on the original trained track (BF = 71.27) and are on average 0.36 cm longer than final path lengths on the trained track (BF = 3.52). For the reverse track, we also find a main effect of set (BFincl > 3 × 10^6^), where initial path lengths in the reverse track are on average 0.80 cm shorter than initial path lengths on the trained track (BF = 121.07) and 0.33 cm longer than final path lengths on the trained track (BF = 3.59). These results show substantial, although incomplete, generalization to the novel track configurations. When restricting the analyses to path lengths inside the track boundaries (Fig. 5C-5D), we do not find evidence for any effects in comparing the different sets of laps. Taken together, these findings show that when it comes to moving in the counterclockwise direction, improvements with training substantially transfer to novel track configurations, but overall movement efficiency is poorer compared to moving in the clockwise direction.

Finally, in both studies we compared path lengths from the last set of laps in the first trained track block with those from the first set of laps in the final trained track block during Session 2 (trained track blocks in Fig. 4B,4D). Participants exhibited longer initial path lengths in the final compared to the first trained track block (Study 1: BF = 131.22; Study 2: BF = 308.75), where path lengths in the final block were 0.56 cm longer on average in Study 1 and 0.50 cm longer in Study 2. Together, these results indicate that interference from training on the novel track configurations has a minimal impact on performance on the original trained track.

### Faster and more efficient movements for clockwise versus counterclockwise movement directions

We then directly compared performance between the two studies using mixed-design ANOVAs, with trial set as a within-subjects factor and study as a between-subjects factor, across all three dependent measures. Although study × trial set interaction effects are reported where applicable, the study-specific analyses presented above already capture within-study performance changes. Accordingly, in this section we focus on the main effect of study to compare overall performance across sessions between the two studies. For lap times, we observe a main effect of study in both sessions (Session 1: BFincl > 4 × 10⁴, with a study × trial set interaction, BFincl = 552.56; Session 2: BFincl = 39.85), indicating that overall movement speed is slower in Study 2, except during the reverse track block (Fig. 2). For accuracy, we also find a main effect of study in both sessions (Session 1: BFincl = 4.07; Session 2: BFincl = 20.18), with slightly higher accuracy in Study 2 than in Study 1, particularly during the final set of Session 1 and the reverse track block in Session 2 (Fig. 3). For movement efficiency, both path length and path length inside the track were generally longer in Study 2 than in Study 1 (path length - Session 1: BFincl > 10⁹; Session 2: BFincl > 3 × 10¹⁰⁷, with a study × trial set interaction, BFincl > 8 × 10⁸⁵; path length inside track - Session 1: BFincl > 2 × 10⁹; Session 2: BFincl > 3 × 10²⁶, with a study × trial set interaction, BFincl > 9 × 10¹⁶; Fig. 4-5). Notably, during the reverse track block, overall path lengths were slightly longer in Study 1 than in Study 2 (Fig. 4B, 4D), whereas path lengths inside the track did not differ between studies for the reverse track (Fig. 5B, 5D). This pattern further highlights reduced movement efficiency in the reverse track block of Study 1. Taken together, our results indicate that movements performed in the counterclockwise direction around the track are overall slower and less efficient than movements performed in the clockwise direction.

## Discussion

We examined practice-related changes in motor execution using a novel gamified two-dimensional racing task, quantifying learning, retention, and generalization with measures commonly used in skill acquisition and adaptation research. Across two studies, movement speed improved rapidly during training. These gains were robustly retained across sessions, although not fully, as initial speed in Session 2 remained slower than the asymptotic level reached at the end of Session 1. Performance transferred substantially to novel track configurations (rotated and reverse tracks), and experience with these tracks did not alter performance when participants returned to the trained track. For accuracy, performance was strongly retained but showed no evidence of offline gains. That is, initial accuracy in Session 2 only matched end-of-training accuracy levels in Session 1. Accuracy also declined transiently when participants first transitioned to novel tracks, suggesting initial interference from the originally trained configuration. However, experience with the novel tracks did not affect subsequent performance on the trained track. Movement efficiency likewise showed strong retention across sessions in both studies. In contrast to speed and accuracy, however, transfer revealed a clear movement direction effect. Movement efficiency partially transferred to the rotated track but not to the reverse track, showing that counterclockwise movements were less efficient than clockwise movements. This asymmetry persisted even when participants trained in the counterclockwise direction during Study 2, with slower speeds and longer paths relative to performance in the clockwise direction. Moreover, counterclockwise training impaired subsequent performance on the clockwise track, where movements would otherwise be expected to be efficient as shown in Study 1. Together, these findings show that while practice yields durable and broadly transferable improvements in motor execution, movement direction imposes a fundamental constraint that shapes how well these improvements generalize to novel contexts.

Our racing task required participants to prioritize both speed and accuracy, allowing us to assess retention and generalization of motor execution improvements. In skill acquisition paradigms, retention is often accompanied by offline gains, such that performance at the start of a second session exceeds the asymptotic level reached at the end of initial training (23). Similar improvements in accuracy across multiple days have been reported in skill-based practice tasks (4,5,7,6,8). In contrast, both our studies show that although accuracy at the start of Session 2 was slightly higher than at the start of Session 1, it did not exceed end-of-training performance from Session 1. Thus, we observed robust retention without evidence of offline gains. One explanation may be ceiling effects, as participants performed with high accuracy from the outset, limiting any further measurable improvements in the task. Moreover, unlike prior studies that constrained movement speed to examine shifts in speed–accuracy tradeoffs, our task encouraged participants to progressively increase speed. This design provides a more complete view of performance retention across multiple behavioral measures, including movement speed and efficiency. Movement speed in Session 2 was slower than asymptotic performance in Session 1 but remained substantially faster than initial performance in Session 1, indicating partial retention. Movement efficiency, indexed by path length, remained stable across sessions. Together, these results show consistent and robust retention across measures, despite a lack of evidence for offline gains.

We also examined whether performance gains generalized to novel track configurations. Prior work shows that improvements acquired within a specific speed range transfer to other speeds (4), but not when movements are performed in a new direction with the same effector or with the untrained effector (7,6). In those cases, apparent improvements are attributed to task re-exposure rather than true transfer (6). Focusing on a single effector, we tested transfer both to a rotated version of the track and to reversed movement direction. Accuracy declined initially when participants encountered either of the novel configurations, suggesting transient interference from the trained track. Interestingly, we observe such interference when switching from the trained track to the novel tracks, but not when participants re-encountered the trained track during the final block in Session 2. This shows how well training improvements are retained and that these learned improvements are robust from interference despite experiencing the novel track configurations. As for measures of speed and movement efficiency, we observe clear transfer for both in the rotated track. Thus, improvements in motor execution acquired in one orientation does substantially generalize to new spatial configurations.

Although performance generalizes to new spatial configurations, movement direction markedly affects transfer. On the reverse track, participants who trained in the clockwise direction showed substantial transfer of movement speed to counterclockwise movements. However, path length measures revealed reduced efficiency for counterclockwise movements. Notably, we also show that this reduced efficiency is independent of accuracy in the task. That is, counterclockwise movements hovered more around the track midline, whereas clockwise movements followed a more optimal trajectory. Study 2 confirmed that this asymmetry reflects effector-specific directional biases. Even when participants trained in the counterclockwise direction, overall speed and efficiency remained worse than in those that trained in the clockwise direction. Moreover, this counterclockwise training impaired subsequent clockwise performance, with movement efficiency remaining impaired despite the optimal clockwise movement performance observed in Study 1. This directional bias may stem from habitual motor experiences in right-handed individuals (e.g. experiences in tasks like writing, (28)). Given existing asymmetries between both dominant and non-dominant effectors (7,6), it remains an open question whether similar movement direction biases in our task will be observed in left-handed people. We also speculate that our findings diverge from prior skill-based practice studies because of fundamental task differences. Unlike Gonda et al. (6), which used discrete point-to-point movements within a semicircular path, our racing task required continuous movement through a curved, elbow-shaped loop, resembling a planned sequence of actions. Consistent with evidence that movement execution differs for single targets versus multi-step sequences (29), the observed direction effect in our task may reflect processes related to planning the full movement around the track (30). Future work, however, should directly test whether planning and execution in our task engage mechanisms similar to those underlying movement sequences.

In conclusion, our findings show that practice refines motor execution such that improvements in speed, accuracy, and efficiency are robustly retained and largely generalize across spatial contexts. At the same time, movement direction constrains this generalization, as clockwise and counterclockwise movements reveal persistent directional biases, likely rooted in effector-specific experience and movement planning demands in our task. By applying measures of learning, retention, and generalization commonly used in adaptation and skill acquisition research, we show that improvements in motor execution are governed by mechanisms closely related to those underlying motor skill acquisition and adaptation.

## Acknowledgments

This work was supported by NSERC for D.Y.P.H.; NSERC - PGSD, OGS, and VISTA scholarships for R.Q.G. The funders had no role in study design, data collection and analysis, decision to publish, or preparation of the manuscript.

## Grants

This study was funded by a NSERC Discovery grant for D.Y.P.H.

## Disclosures

The authors declare no competing financial interests.

## Author contributions

D.Y.P.H. designed the research. S.N. and R.Q.G. collected the data. R.Q.G. and A.K. contributed experimental and analytic code. S.N. and R.Q.G. analyzed the data. R.Q.G. wrote the main manuscript text, which was carefully edited by all authors.

## Notes

https://doi.org/10.17605/OSF.IO/W5D4U

### Competing Interest Statement

The authors have declared no competing interest.

